# Nectar compounds impact bacterial and fungal growth and shift community dynamics in a nectar analog

**DOI:** 10.1101/2022.03.29.485809

**Authors:** Tobias G. Mueller, Jacob S. Francis, Rachel L. Vannette

## Abstract

Floral nectar is frequently colonized by fungi and bacteria. However, within individual flowers, nectar microbial communities are typically species-poor and dominated by few cosmopolitan genera. One hypothesis is that nectar constituents may act as a strong environmental filter. Non-sugar constituents in nectar could affect species composition via broad antimicrobial activity or differential effects on nectar microbial species. Here, we tested how five non-sugar nectar compounds as well as elevated sugar impacted the growth of 12 fungal and bacterial species isolated from flowers, pollinators, and the environment. We hypothesized that microbes isolated from nectar would be better able to grow in the presence of these compounds. Additionally, to test if nectar compounds could affect the outcome of competition among microbial taxa, we grew a subset of microbes in co-culture assays across a subset of treatments.

We found that some compounds such as H_2_O_2_ broadly suppressed microbial growth across many but not all microbes tested. Other tested compounds were more specialized in the microbes they impacted. As hypothesized, the nectar specialist *Metschnikowia reukaufii* was unaffected by most nectar compounds assayed. However, many non-nectar specialist microbes remained unaffected by compounds thought to reduce microbial growth in nectar. Our results show that nectar chemistry can influence nectar microbial communities but that microbe-specific responses to nectar compounds are common. Nectar chemistry also affected the outcome of species interactions among microbial taxa, suggesting that non-sugar compounds in nectar can affect microbial community assembly and abundance in flowers.

## Introduction

Most angiosperms produce floral nectar to attract pollinators. Floral nectar (hereafter simply nectar) is an aqueous solution often predominantly composed of sugars including sucrose, glucose, and fructose [1]. However, nectar is much more than a simple sugar solution; approximately 10% of nectar’s dry weight is composed of non-sugar compounds including free amino acids, proteins, lipids, vitamins, and alkaloids among other compounds [2–4] and can differ substantially among and within species [3, 5].

Nectar can be colonized by microbes, primarily yeasts and bacteria, which are deposited by floral visitors [6–8]. Surveys typically find 20 - 50% of flowers contain culturable microbes depending on plant species and environment [9–13]. The microbes found in nectar can range from plant and pollinator pathogens, to putatively mutualistic fungi, to microbes that may be commensal or have no documented effects on plants or pollinators [14]. Once deposited, nectar microbes can reach high densities, growing to more than 10^5^ cells/μL for yeasts and 10^7^ cells/μL for bacteria [15]. However, microbial communities often exhibit low alpha diversity within individual nectar samples, consisting of a few globally dominant genera, including fungi, such as *Metschnikowia* and *Aureobasidium* [9, 10, 16], and bacteria, such as *Acinetobacter* [12, 17–19]. The microbes that establish in nectar are a subset of the microbes carried by pollinators and in the environment [17, 20, 21]. While it is clear that many microbes deposited in floral nectar fail to establish [20–22], numerous processes may generate the low microbial diversity observed in nectar. Possible mechanisms include differential dispersal of microbes [7]; competitive exclusion that favors early arriving, faster growing, or inhibiting species [23, 24]; or strong filtering by the chemistry of the nectar environment [21]. These mechanisms are not mutually exclusive and likely vary in importance depending upon context. However, in some systems animal-flower visitation networks alone cannot explain nectar microbial communities suggesting that filtering may play a role [7].

Some nectar traits are thought to provide antimicrobial activity [21, 25]. The high sugar concentrations in nectar leads to extreme osmotic pressure and high C:N ratios both of which limit microbial growth [21, 26, 27]. Additionally, antimicrobial compounds are commonly produced in nectar [25, 28]. In ornamental tobacco (*Nicotiana langsdorffii × Nicotiana sanderae*), hydrogen peroxide levels can reach 4mM [29], suppressing some but not all microbes’ growth [30]. Other antimicrobial proteins are thought to have activity against specific groups of microbes [25]. In previous comparative studies, nectar compounds including hydrogen peroxide, the antimicrobial protein BrLTP2.1, and the floral volatile linalool showed species-specific effects, reducing microbial growth for some species but not others [28, 30–32]. However, few studies have broadly compared if microbes, isolated from nectar and other habitats, vary in resistance to a range of nectar compounds (however, see [20, 32, 33]), and if these compounds impact microbe-microbe interactions.

Here, we use *in-vitro* growth assays to test the degree to which nectar chemistry alone, or in combination with competitive dynamics, impacts microbial growth in a nectar analog. First, we tested the hypothesis that common nectar microbes can better tolerate a variety of nectar chemistries compared to microbes isolated from non-nectar habitats. If non-nectar specialists grow well in the presence of nectar compounds, it would indicate that filtering by these compounds is not a major driver of community assembly, and that other factors such as dispersal limitation or competition are more important. However, if only nectar specialists can maintain growth in the presence of common compounds found in nectar, it would suggest that environmental filtering may play a major role in nectar microbial community assembly. Second, we tested the hypothesis that the presence of nectar compounds affects the outcomes of microbial competition in nectar.

## Methods

### Microbial strains

We tested the effects of nectar compounds on the growth of the yeasts *Metschnikowia reukaufii, Aureobasidium pullulans, Starmerella bombi, Rhodotorula fujisanensis, Saccharomyces cerevisiae, Zygosaccharomyces bailii*, and the bacteria, *Acinetobacter nectaris, Rosenbergiella nectarea, Bacillus subtilis, Pantoea agglomerans, Pseudomonas mandelii, Pectobacterium carotovorum*. The species assayed include microbes commonly isolated from nectar, pollinators, and the environment (Table 1).

**Table 1.**
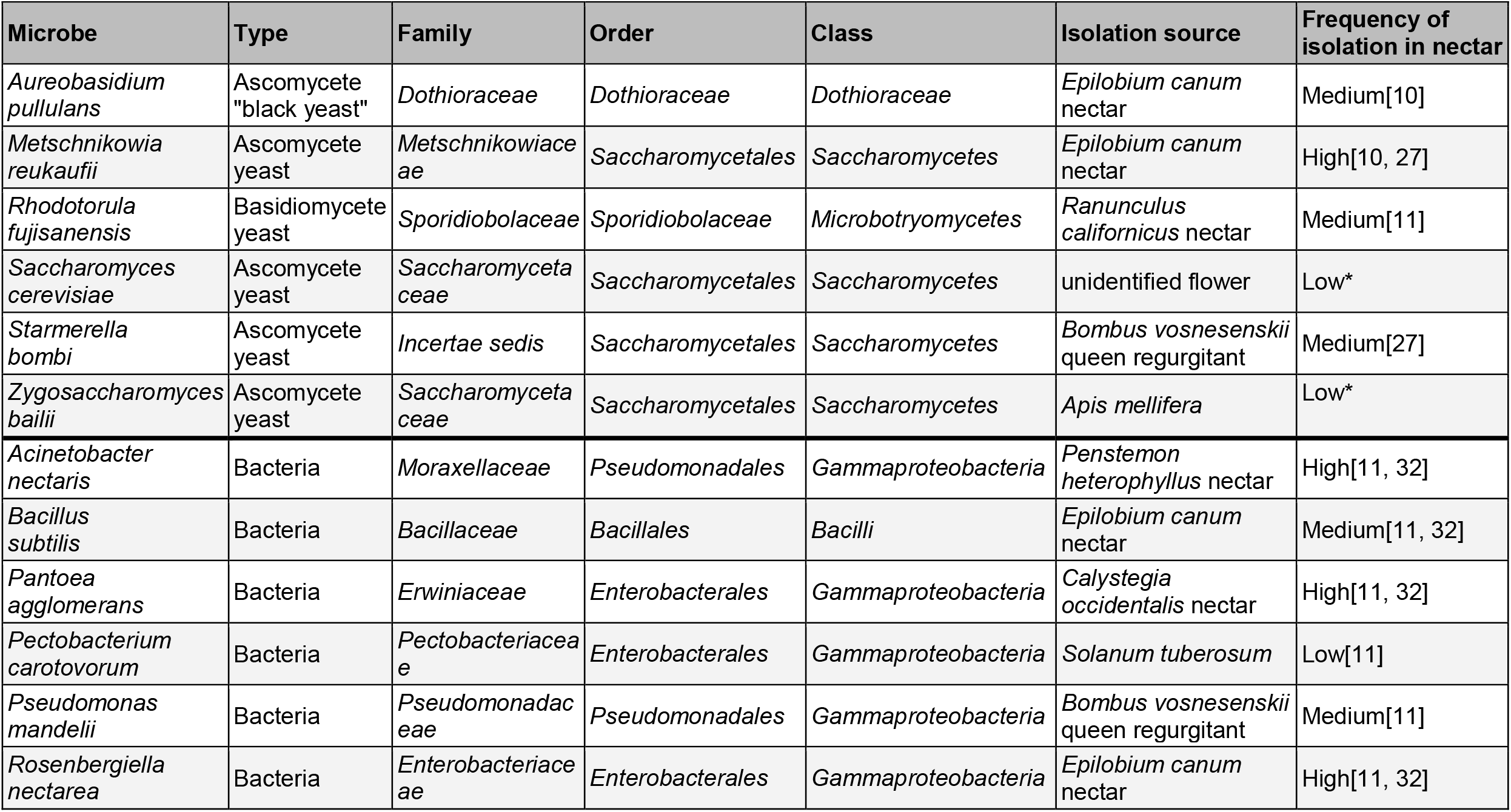
The microbes used in the study along with the strain’s source. The prevalence score is an approximation based on the frequency microbes have been discovered in nectar microbe surveys. * indicates we are not aware of this species being documented as isolated from floral nectar

To standardize microbial density across treatments, we created master microbial suspensions. First, we grew microbial stocks for three days on Yeast Media Agar (YMA) containing chloramphenicol (at 100 mg/L to reduce bacterial growth) or fructose enhanced Trypticase Soy Agar (TSA) containing cycloheximide (at 100 mg/L to reduce fungal growth) for fungi and bacteria respectively (Supplemental Method 1). We then created 2000 cell/μl suspensions (quantified via hemocytometer) in 15% glycerol and stored aliquots at -80^°^C for the duration of the experiment. Each plate or cogrowth assay used a new aliquot to ensure that every replicate had the same starting cell densities of each focal microbe.

### Chemical constituents

We tested compounds detected in nectar that have been hypothesized or demonstrated to be antimicrobial and used concentrations previously documented in nectar (Supplemental Table 1). We tested hydrogen peroxide (H_2_O_2)_, a reactive oxygen species found in some nectars at two concentrations (2mM and 4mM, [29]); deltaline, a norditerpene alkaloid found in the nectar of *Delphinium spp*. and a potent toxin for eukaryotes (22ug/ml, [34]); BrLTP2.1, a lipid transfer protein isolated from *Brassica rapa* nectar, hereafter referred to as LTP (150μg/ml, [28]); linalool, a common volatile found in nectar (100ng/ml, [32]); ethanol (EtOH), a common byproduct of fermentation in nectar (1%, [35]) and elevated sugar at 30%, along with a 15% base control nectar solution (which covers the low and moderate levels of natural sugar concentrations) [3]. See Supplemental Table 2 for the recipes of control and treatment “nectars”.

### Preparing synthetic nectars

To prepare the synthetic nectars (treatment and control solutions), we weighed dry reagents on a microbalance to a precision of 0.0025 grams before washing them into a volumetric flask and dissolving the reagents in DI water. Liquid reagents were added and then the entire solution was diluted with DI water to the proper concentration before being vortexed and sterilized using a syringe filter (0.2 μm cellulose acetate membrane, Corning, Corning NY, product number 431219). Base nectar consisted of 15% sugar (50:25:25 sucrose:glucose:fructose) w/v, 1% peptone w/v, 3% yeast extract w/v, 50% 100x non-essential amino acids v/v.

### Plate reader growth assay

To test the effect of individual compounds on the growth of single microbe species, we used 96 well plate growth assays and synthetic nectars. Each well in a plate contained 190μL of treatment or control nectar and 10μL of microbial freezer stock solutions (2000 cells/μL in 15% glycerol v/v with 15% sucrose w/v). Each plate consisted of a single chemical treatment assayed across all 12 microbes (6 treatment and 2 control wells per microbe, see Supplemental Figure 1 for plate mapping). We assigned each microbe’s location on the 96 well plate using a random number generator and kept the location consistent across all plates. After mixing chemical treatments and microbial strains, we triple parafilmed the 96 well plate lid and put it immediately into an optical reader (Biotek synergy HTX, Agilent, Santa Clara CA, USA) which incubated the plate at 30°C, provided continuous linear shaking at 567cpm (3mm), and took optical density measurements at 600 nm every 15 minutes for 72 hours. After preparing each plate, we assessed potential contamination by plating out the control and treatment nectar solutions onto YM and TSA plates. None of our uninoculated synthetic nectars contained culturable microbes.

### Co-growth experiment

To test if nectar composition could shift microbial interactions, we grew pairs of microbes across several treatment solutions. We chose a subset of microbes that produce distinguishable colonies on plates to test the following combinations: 1) a facultative nectar yeast with a non-nectar yeast (*Starmerella bombi & Zygosaccharomyces bailii)*, 2) a nectar specialist yeast with a nectar specialist bacteria (*Metschnikowia reukaufii & Rosenbergiella nectarea)*, and 3) a non-nectar specialist yeast with a nectar specialist bacteria (*Saccharomyces cerevisiae & Rosenbergiella nectarea)*. We also ran a pairing of *Metschnikowia reukaufii & Saccharomyces cerevisiae*, however, the vials exploded during incubation due to extremely rapid fermentation. If the dominance of nectar specialists is driven by nectar chemicals shifting microbe-microbe competition we predict nectar specialists will increase in relative abundance in the presence of nectar compounds, while the relative performance of environmental microbes should be reduced compared to control co-growth trials.

We chose a subset of treatments for co-growth assays, including 4mM H_2_O_2_, 22μg/ml deltaline, 100ng/ml linalool, and 1% EtOH. We did not include LTP due to a limited amount of protein available for assays and did not include 30% sucrose as it showed no significant impacts on growth during our plate reader assays. Treatments used the same recipes as the growth experiments described above.

We performed the co-growth experiment in 200μl 8-strip PCR tubes with 6 tubes per treatment– microbe combination. Each tube consisted of 190μl of synthetic nectar and 5μl (10,000 cells) of each microbial freezer stock in that pairing. We vortexed the tubes for 5 seconds and incubated them at 25°C for 72 hours. To assess the effect of co-growth, we also grew each microbe in isolation, following the same methods, with tubes consisting of 190μL of synthetic nectar and 5ul of microbial suspension (N=4).

After 72 hours of incubation, we serially diluted the microbial suspensions and plated 100μL of diluted microbial suspension onto TSA and YMA plates. We diluted TSA plates 2x (plating 50μl of original suspension), YMA plates from the *Starmerella bombi* and *Zygosaccharomyces* pairing both 20x (5μl plated) and 200x (0.5μl plated), and YMA plates with *Metschnikowia* or *Saccharomyces* 200x (0.5μl plated). We chose these dilutions as they created countable CFUs. We then incubated plates at 25°C for 72 hours to allow microbial colonies to form, after which we counted the number of colonies per plate. *Rosenbergiella* did not form single colonies and instead the percent of the plate covered by growth was estimated and adjusted relative to the maximum CFU count of its yeast pairing (100% coverage = maximum CFU count).

## Analysis

All analyses were performed in RStudio using R version 4.0.1 [36].

### Curve fitting and data curation

We used the Grofit package [37] to fit logarithmic curves to the optical density (OD) timeseries. The initial OD value for each well was deducted from all readings to account for starting solution OD. Best-fit growth curve models were selected using AIC and each fitted curve was visually inspected after which growth rate (**µ**) and maximum OD (**A**) were extracted from the fitted curves.

To ensure data quality, we performed the following checks on all growth curves before curve fitting: if a well did not change OD over 72 hours, the **A** and **µ** were set to zero. If the OD increased and then returned to the starting value within 72 hours, we considered the well having no growth and set both parameters to zero. If only 1 treatment well out of the 6 did not grow, we considered this to be due to an error (possibly no microbe addition) and removed the well from the analysis. If no mathematical fit could be plotted to the OD readings, we removed the well from the analysis (less than 5% of growth curves). Many of these unfittable wells showed flatline curves with a single value change during the 72 hours likely not caused by microbial growth. One well showed an **A** 100 times greater than all other wells in that treatment and was removed. For a single plate (deltaline), condensation caused a temporary drop in the OD for the first 45 minutes. For these wells we set the starting OD as the lowest OD reading from the first hour.

To control for variation in growth among plates, we divided the mean growth of control wells for each microbe on each plate by the mean growth of that microbe’s control wells across all plates giving us a plate-specific growth ratio. We then multiplied the treatment wells on a given plate by that plate-specific ratio. To compare a treatment’s relative impact on growth across microbes, we scaled all microbes’ **µ** and **A**. This was done by adjusting growth relative to each microbe’s growth in control nectar across all plates *(scaled value = treatment* **µ** or **A** */ mean control* **µ** or **A**). A scaled value over 1 indicates a treatment **µ** or **A** greater than that microbe’s control and scaled value below 1 indicates a **µ** or **A** lower than that microbe’s control. These transformations allow us to compare the effects of nectar compounds across many microbes that varied in absolute growth.

### Treatment impacts across all microbes

To compare the effects of treatment across all microbes we fit a negative binomial model [38] with scaled maximum OD as a function of treatment. To test if the scaled maximum OD and scaled growth rate were correlated we calculated the Pearson’s correlation coefficient.

### Microbe-specific response to treatments

We compared each microbe’s growth in different nectar chemistries to their growth in control nectar using a Kruskal-Wallis test followed by a Dunnett’s test [39], with separate models for maximum OD and maximum growth rate. To test if treatment impacts were related to the frequency microbes occur in nectar (hereafter referred to as nectar specialization) we ranked microbes as “high”, “medium”, or “low” according to their relative incidence and abundance in nectar (Table 1). We ran a Kruskal-Wallis test comparing scaled maximum OD and scaled growth rate to the level of specialization. Significant results were followed by a Dunn’s test with a Holm-Bonferroni correction to compute pairwise differences [39].

### Differences between yeast and bacteria

To compare if nectar chemistry differentially affected yeasts compared to bacteria, we used a linear random effects model (for maximum OD) and negative binomial model (for scaled maximum OD) comparing microbial growth between kingdoms, with microbe and treatment as random intercept terms.

### Co-growth assay

To compare how nectar chemistry can change community dynamics, we used a Kruskal-Wallis test followed by a Dunn’s test comparing each microbe’s growth in co-culture across different nectar chemistries and alone in control nectar.

## Results

### Treatment impacts across all microbes

Nectar compounds differed in their effect on maximum scaled OD (Figure 1); H_2_O_2_ strongly suppressed the growth of most microbes at 2mM (linear mixed model coefficients and standard error: -1.03 +/- 0.24, p < 0.001) and 4mM (−2.01 +/- 0.33, p < 0.001). 30% sucrose (−0.007 +/- 0.15, p = 0.65), LTP (−0.11 +/- 0.147, p = 0.47), and linalool (−0.14 +/- 0.15, p = 0.37) showed a (non-significant) trend of maximum OD suppression overall. In contrast, the diterpene alkaloid deltaline increased maximum OD overall (0.6 +/- 0.12, p < 0.001) and EtOH had no significant effect (0.2 +/- 0.14, p = 0.15). Scaled maximum OD was correlated with scaled maximum growth rate (r = 0.53, p < 0.001) and effects of treatments on both were congruent, although not identical (Supplemental Figure 2).

**Fig. 1.**
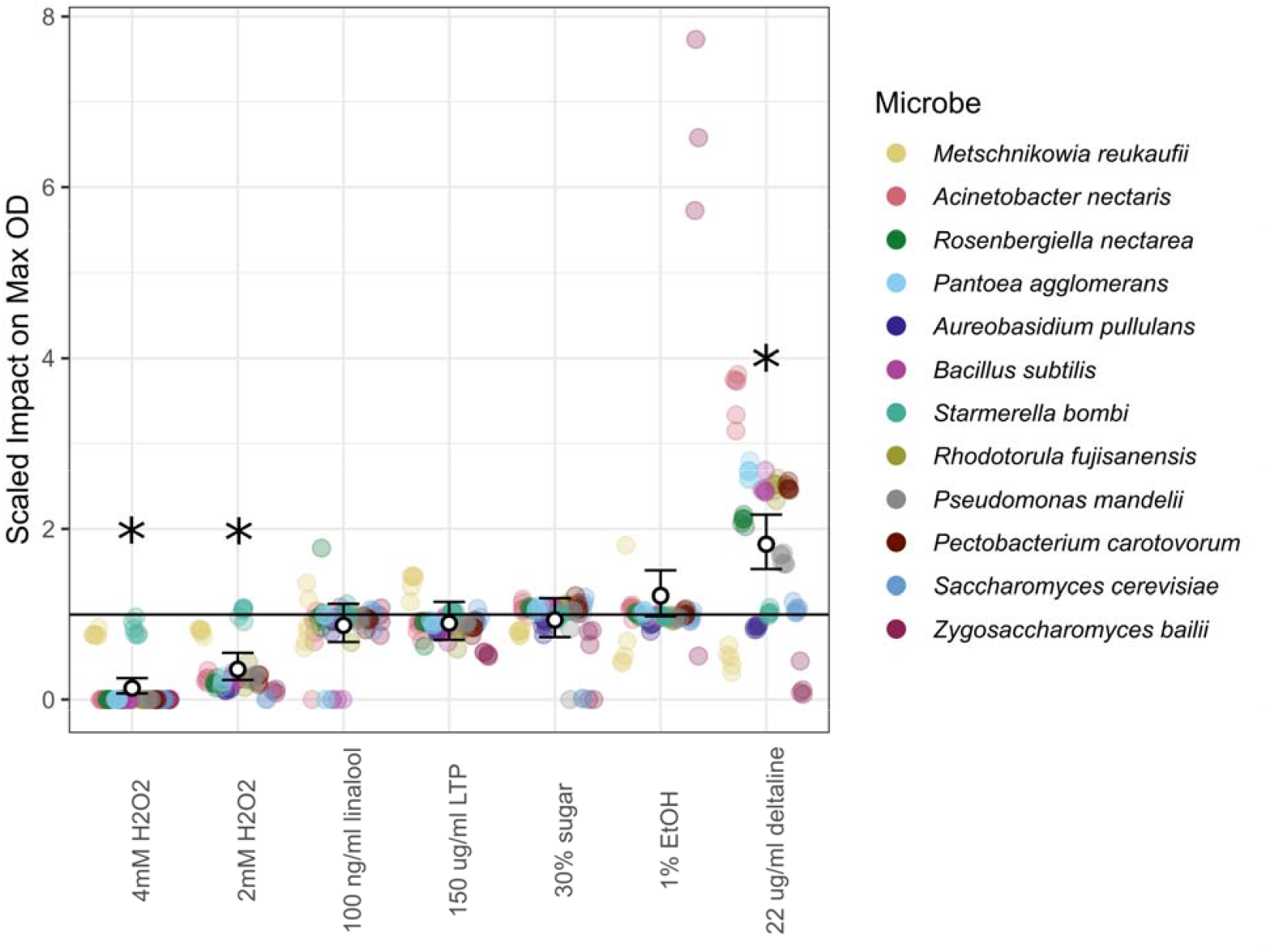
Nectar compounds differ in their effects on maximum microbial density. The Y-axis indicates the scaled effect of treatment on maximum OD (optical density) compared to control nectar. A value of 1 represents equal density in treatment and controls; values higher than 1 represent an increase in maximum density compared to controls and values lower than 1 indicate a decrease in maximum density. White points and bars show the negative binomial model coefficient and 95% confidence intervals for each compound. Colored points indicate individual replicates for each microbe and contain a slight horizontal jitter to aid in readability. A horizontal line is added at Y=1. Stars represent significant overall treatment impacts at p < 0.05

### Microbe-specific response to treatments

Microbial species varied in their maximum OD and growth rate in control conditions and in response to treatments (Supplemental Figures 3-4, p < 0.05). All microbes were impacted by at least one treatment, but treatments differed in their effect on maximum OD (Figure 2) and growth rate (Figure 3) across microbial species. Species’ responses to nectar composition depended on the specific nectar compound tested: no microbe had significantly reduced maximum OD or growth rate across all treatments (Figure 2). When comparing across all treatments, the scaled maximum OD was not significantly different across degrees of nectar specialization (p > 0.05; Figure 4a), however, scaled growth rate was significantly different: microbes infrequently isolated from nectar had a lower scaled growth rate than both the highly and medium specialized group (p < 0.05; Figure 4b).

**Fig. 2.**
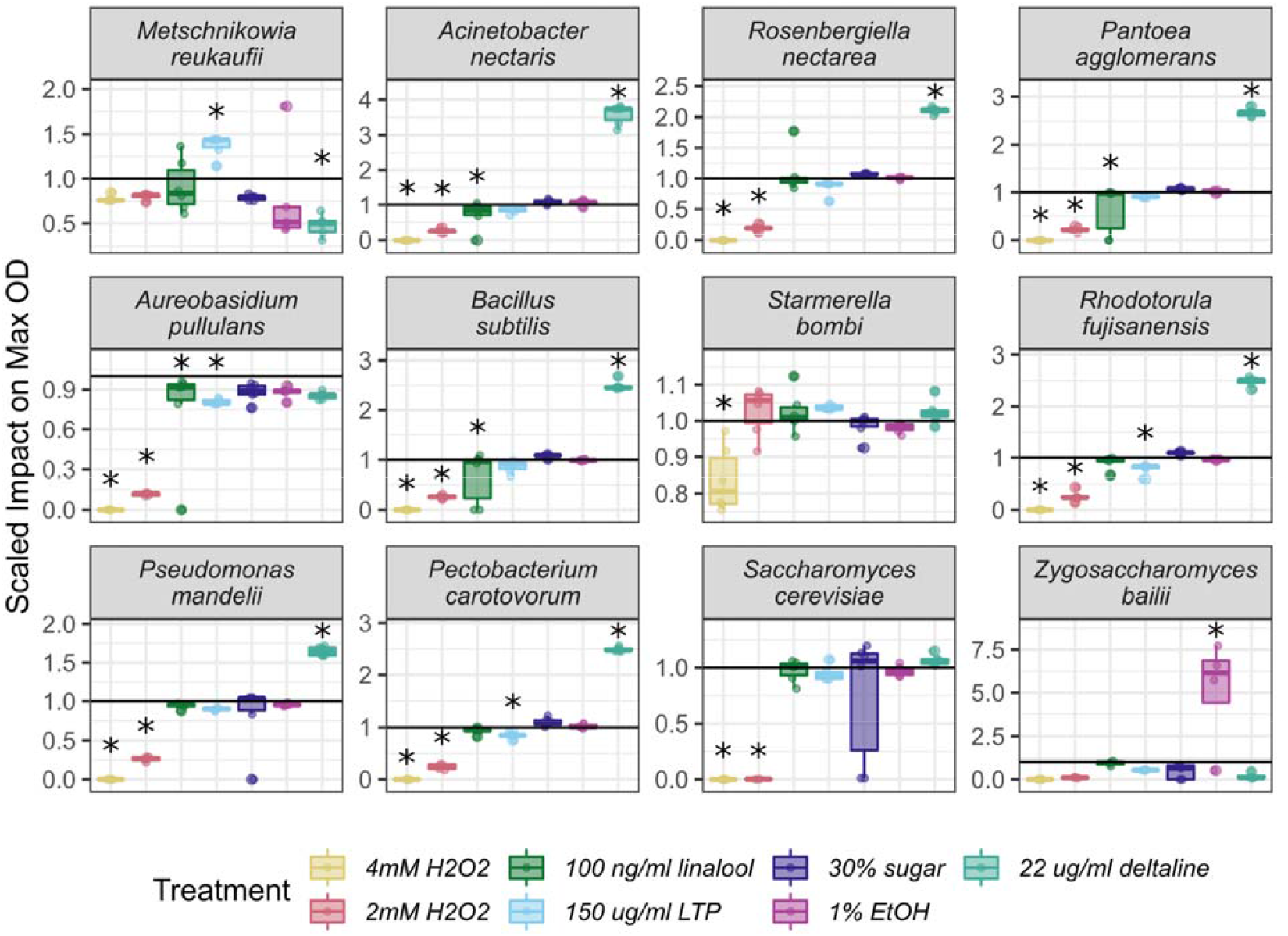
Microbial species vary in the scaled impact of treatment on maximum optical density. The Y axis is the scaled impact of a treatment on a microbe’s maximum OD compared to controls, as in Fig 1, but separated to more clearly display variation among species. Microbes are ordered from most frequently (top left) to least frequently isolated from nectar (bottom right). Stars indicate significant treatment impact on maximum OD compared to control (p < .05). See Supplemental Figure 3 for non-scaled data

**Fig. 3.**
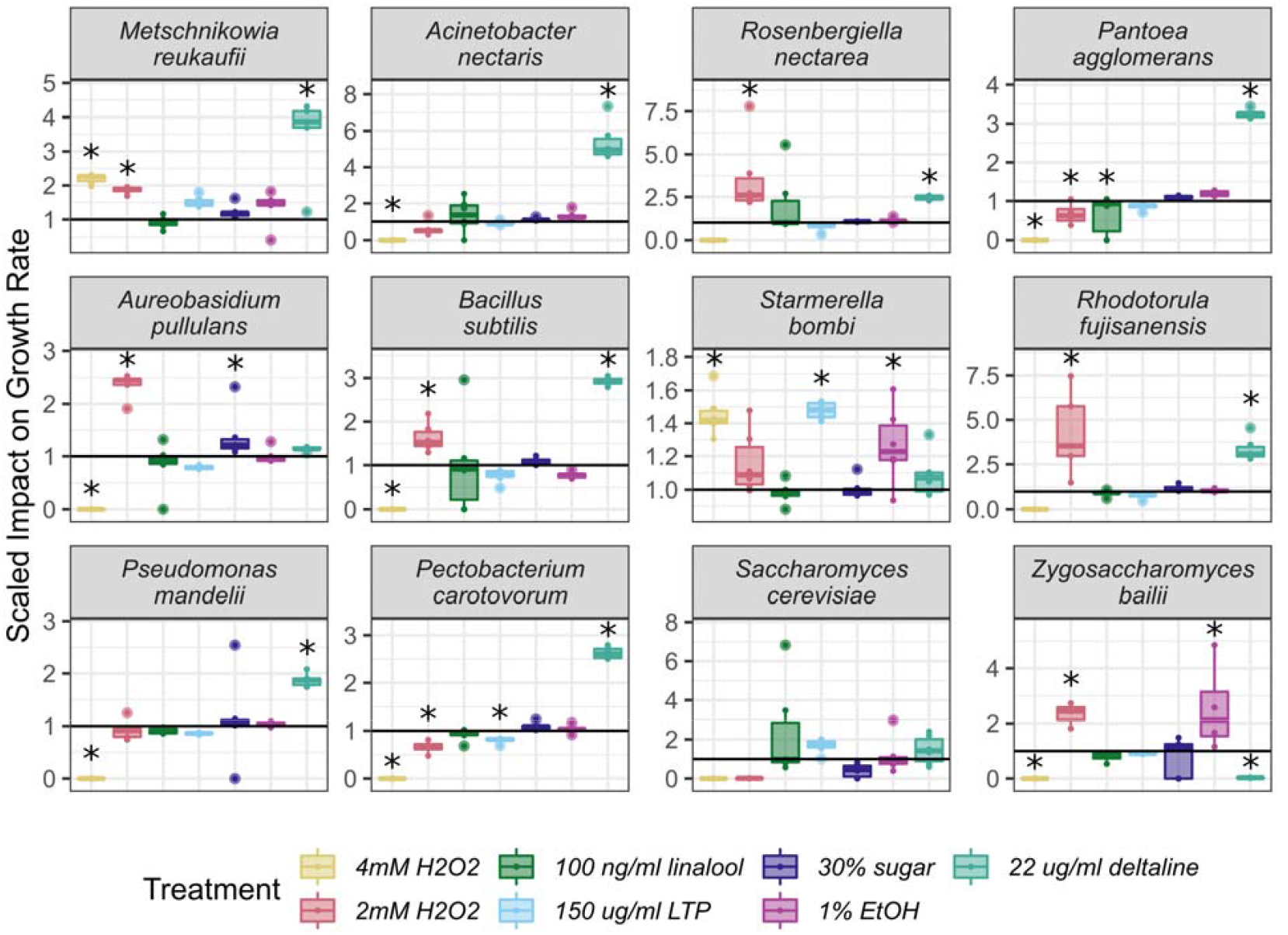
Microbial species vary in the scaled impact of treatment on growth rate. The Y axis is the scaled impact of a treatment on a microbe’s growth rate compared to controls, as in Fig 1, but separated to more clearly display variation among species. Microbes are ordered from most frequently (top left) to least frequently isolated from nectar (bottom right). Stars indicate significant treatment impact on maximum OD compared to control (p < .05). See Supplemental Figure 4 for non-scaled data

**Fig. 4.**
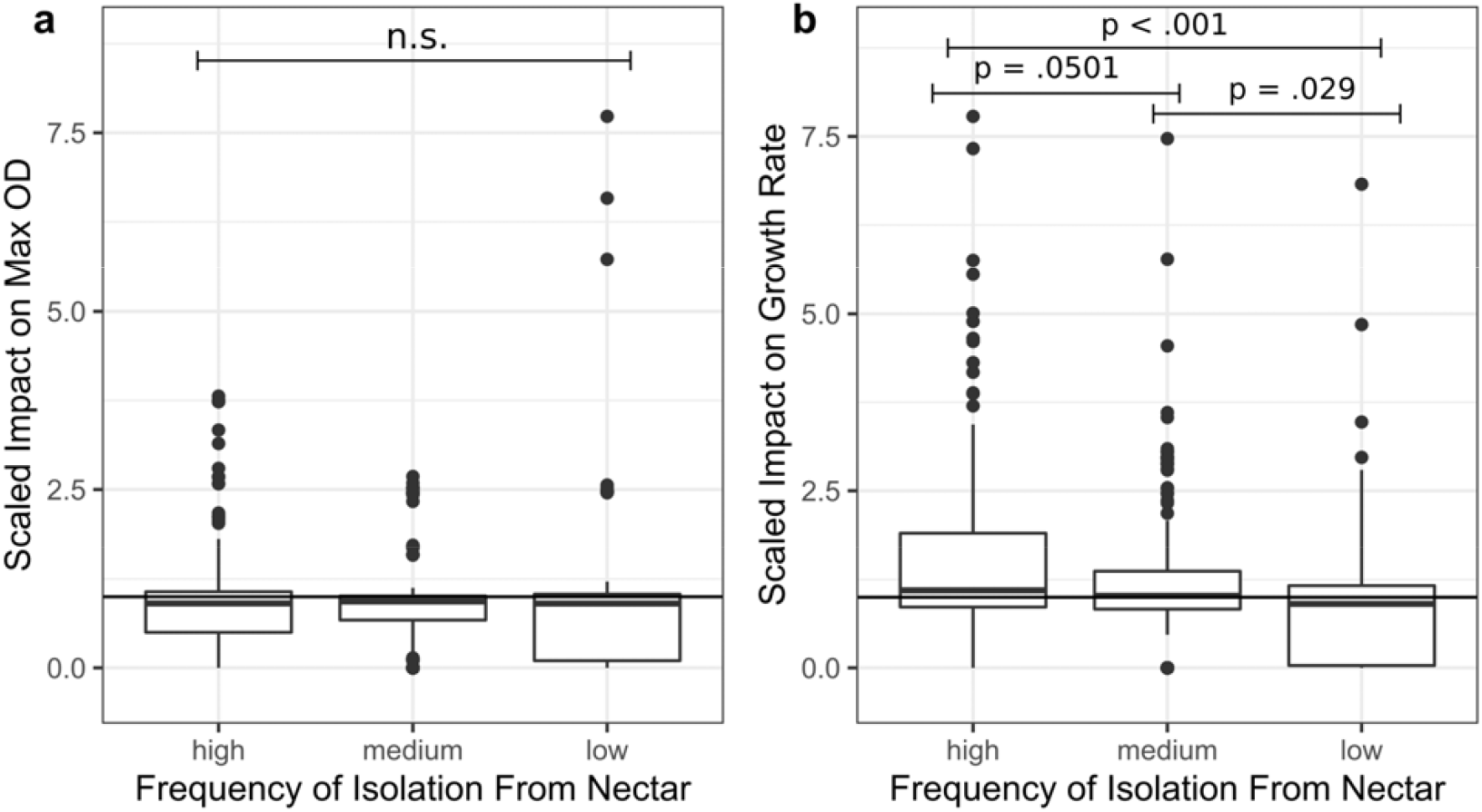
Microbial isolation source predicts sensitivity of growth rate but not maximum OD to treatments. The Y-axis indicates the scaled effect of treatment on maximum OD (optical density; panel a) and growth rate (panel b) compared to control nectar. A value of 1 represents equal density in treatment and controls; values higher than one represent an increase in maximum density compared to controls and values lower than one indicate a decrease in maximum density. A horizontal line is added at Y = 1

### Differences between yeast and bacteria

Yeasts and bacteria differed significantly in the maximum OD attained, with yeasts (0.82 +/- 0.35, p = 0.04) having a higher max OD than bacteria (0.01 +/- 0.25, p = 0.96) (Supplemental Figure 5a). However, when assaying treatments’ scaled impact on max OD there was no significant difference between yeasts (−0.09 +/- 0.09, p = 0.33) and bacteria (−0.3 +/- 0.26, p = 0.25) (Supplemental Figure 5b), indicating that while yeasts generally grow to optically denser levels in synthetic nectar they are not more or less resistant to the chemicals assayed compared to bacteria.

**Fig. 5.**
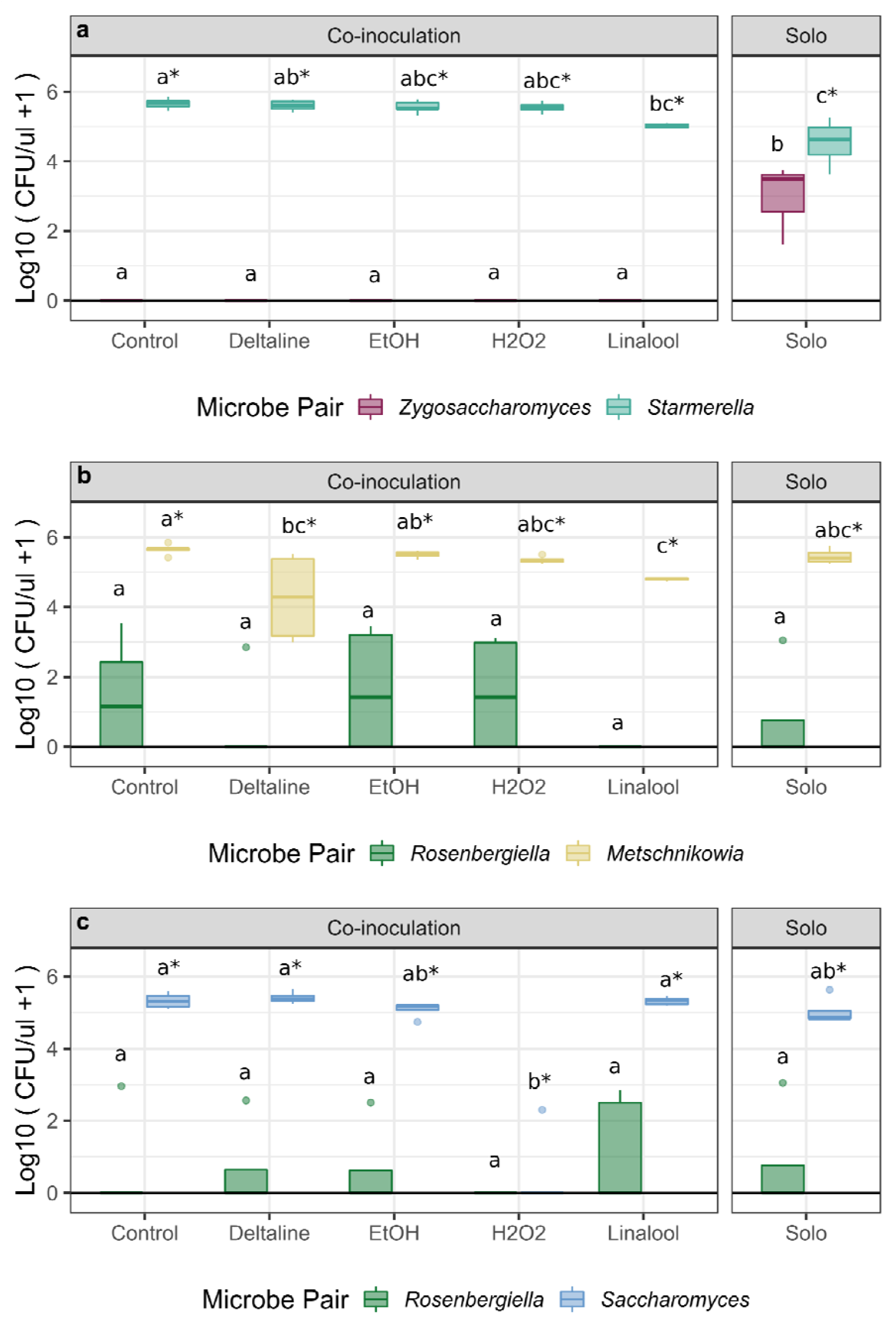
Nectar compounds influence microbial community outcomes but differ depending on species considered. The colony forming units (CFUs) per ul of synthetic nectar formed by microbes grown in co-culture and alone across different nectar chemistries. Each panel represents a different pairing of microbes; panel A pairs a facultative nectar yeast with a non-nectar yeast (*Starmerella bombi & Zygosaccharomyces bailii)*, panel B pairs a nectar specialist yeast with a nectar specialist bacteria (*Metschnikowia reukaufii & Rosenbergiella nectarea)*, and panel C pairs a non-nectar specialist yeast with a nectar specialist bacteria (*Saccharomyces cerevisiae & Rosenbergiella nectarea)*. Letters indicate significant pairwise differences between treatments (p < .05) and are shown separately for each microbe

### Co-growth assay

The presence of competitors and nectar compounds together affected microbial abundance after 3 days for all species pairings (Figure 5). For example, in a co-culture of the food spoilage specialist *Z. bailii* and bee-associated *S. bombi, Z. bailii* never formed CFUs in the presence of a competitor, but did when grown alone, suggesting strong competitive exclusion. In contrast, *S. bombi* in the same pairing showed increased CFU formation in co-culture relative to its growth alone (p < 0.001), in control nectar, 22ug/ml deltaline, 1% EtOH, and 4mM H_2_O_2_ treatment nectars (Figure 5a). In the pairing of two nectar ‘specialists’, neither the bacteria *R. nectarea* nor the yeast *M. reukaufii* showed an altered CFU density in co-culture compared to growth in isolation (Figure 5b). When co-culturing *R. nectarea* and *S. cerevisiae*, we found that contrary to our original hypothesis, the non-nectar yeast *S. cerevisiae* did not show a significant reduction (p > 0.05) in growth compared with growth alone. Notably, however, the addition of H_2_O_2_ reduced *S. cerevisiae* and made *R. nectarea* growth undetectable (Figure 5c) — in contrast to the ability of *R. nectarea* to persist in the presence of *M. reukaufii* in H_2_O_2_-containing nectar.

## Discussion

All nectar constituents tested had species-specific effects on microbial growth, significantly impacting certain microbes while showing no impact on others. Hydrogen peroxide showed strong antimicrobial properties across most microbes assayed, both nectar specialists and non-specialists. It is unknown how common H_2_O_2_ is in nectar, but it has been detected in several genera of plants including *Nicotiana* and *Cucurbita* [30, 40]. Despite strong suppressive effects on most species (including those with documented catalase activity) [12], the antimicrobial effect of H_2_O_2_ was not universal. Notably the maximum OD of the yeast *M. reukaufii* and *Zygosaccharomyces bailli* were unaffected by any concentration of H_2_O_2_ tested and *S. bombi* was only affected at 4mM. Other tested compounds were more selective in their growth suppression and impacted different microbes including those frequently and seldom isolated from nectar.

The observed differences in the selectivity of compounds suggest that nectar antimicrobial compounds (NACs) may fall into two broad classes with different functions: general antimicrobials and selective filters. General NACs (e.g., H_2_O_2_ here) may keep a flower from being colonized by most microbes and are possibly common in nature. In some ecosystems as many as 80% of plants have no culturable yeasts and some have very low incidence of culturable bacteria [13, 41]. We predict that general NACs, or other mechanisms to limit microbial growth, might be more common in ecosystems where plants have a high likelihood of colonization by antagonistic microbes but a low probability of colonization by beneficial microbes (or where the costs of antagonists consistently outweigh the benefits of mutualists). Conversely, we predict that selective filtering NACs might be more common in ecosystems where plants have equal likelihoods of being colonized by beneficial or antagonistic microbes. Direct effects of NACs on pollinator behavior and health, however, should not be discounted and likely also plays a role in the selection on NACs [42]. While we lack data on the plant traits that shape communities of antagonistic and beneficial microbes [14], and there are likely other modes beyond NACs that work in conjunction such as floral morphology or other nectar constituents including enzymes, ions, lipids, among others, these data suggest that selective NACs may be one route by which plants shape their nectar microbiome. However, with the extreme diversity in floral nectar chemistry, many general and selective NACs have likely not yet been identified or may escape notice by being context dependent. Characterizing the relative abundance of general and selective NACs across different microbial landscapes might be particularly fruitful in disentangling how microbes shape selection on nectar traits.

Our findings suggest that NACs can also shift competitive dynamics and the trajectories of nectar microbial communities as previously suggested [15]. While we found no relationship between degree of nectar specialization and treatment impacts on maximum growth, the growth rate of non-nectar specialists was more suppressed in the presence of nectar compounds, which could affect community assembly. Our co-culture experiment further shows that treatments can impact communities not only by decreasing the growth of some microbes, but also increasing the growth of others in co-culture. Here, *Z. bailii* did not grow in co-cultures with *S. bombi*, however, *S. bombi* showed elevated growth in co-culture, even in the presence of H_2_O_2_. We hypothesize that the presence of *Z. bailii* may have facilitated the growth of *S. bombi* by potentially providing additional nutrition. Alternatively, it appears that some microbes may facilitate each other’s growth. For example, *R. nectarea* grew in H_2_O_2_-containing nectar in the presence of *M. reukaufii* but not *S. cerevisiae*, perhaps suggesting that *M. reukaufii*, which itself does not appear to be impacted by H_2_O_2_, may have methods for detoxifying H_2_O_2_ that extend to other inhabitants of the same nectar environment.

The impact of plant chemistry on ecological interactions can be difficult to predict and some presumptive NACs may even benefit certain microbes. We predicted that the norditerpene alkaloid deltaline would broadly suppress microbial growth, but our results generally suggest otherwise. Deltaline only decreased the growth of *M. reukaufii*, with most other microbes increasing in maximum OD relative to their control. This is surprising considering that other norditerpene alkaloids, extracted from flowering plants in the same family as *Delphinium*, have strong antimicrobial properties [43]. Prior work looking at the antimicrobial effects of norditerpenes, however, tested concentrations higher than those occurring in nectar [43]. For microbes that do not experience growth suppression, it is possible that deltaline is a source of otherwise limiting nitrogen [44]. This might be particularly true in nectar (or nectar surrogates). Compounds that might be otherwise anti-microbial in growth media or in other plant tissues may benefit microbes in nectar. These findings highlight that generalizing across plant tissues and among whole classes, or even subclasses, of compounds should be done with caution.

Although the impact of nectar secondary metabolites on microbes may be an understudied ecological role, other abiotic and biotic ecological drivers should also be considered. Nectar chemicals are widespread [45] but may be non-adaptive consequences of chemical defense in other plant tissues [45, 46] where they can effect florivores or pollinators and their behavior [47]. Additionally, nectar chemicals are often in low concentrations when compared to compounds in other plant tissues [48]. Compounds in other plant tissues may also influence the nectar environment and shape microbial communities, for instance, when pollen gets deposited into floral nectar. Nectar is a complex and dynamic solution, changing with enzyme activity, host-mediated secretion and resorption, and via contact with floral tissues – all precluded by our use of synthetic nectar. It is possible that these complex interactions of chemicals may increase or decrease the effect of the specific compounds tested here. Whether the impacts of NACs observed here are stronger or weaker than these other factors (and thus are ecologically relevant) is an open question.

Taken together, our results suggest variable effects of nectar chemistry and that different microbes may be excluded from nectar for varying reasons. The findings that nectar compounds can shift microbial colonization and community dynamics raise more questions for further study. Given that nectar is chemically diverse [48], and microbes vary in dispersal limitation [13], what does the observed selectivity of NACs mean at a landscape scale? On one hand, it could lead to a diversity of microbial niches where different floral species have different selective NACs, and thus floral diversity would likely increase microbial diversity at the landscape scale. However, this is not found in nectar surveys, suggesting that other strong drivers, such as dispersal [8, 13], competitive ability [23, 24], or intraspecific variation in microbial sensitivity to NACs, also contribute to low species diversity in floral microbial communities[49, 50]. Finally, given our result that nectar secondary chemistry can affect microbial growth, characterizing variation in antimicrobial potential among plant populations and species may allow a better understanding of how microbes, pollinators and other forces shape the ecology and evolution of nectar traits.

## Acknowledgements

We especially thank Anthony Schmitt and Clay Carter for isolating and providing the BrLTP2.1 protein and for their early comments on experimental direction. We would like to thank members of the Vannette lab including Shawn Christensen, Amber Crowley-Gall, Marshall McMunn, and Danielle Rutkowski, as well as Kate LeCroy and Scott McArt for their feedback and comments on the manuscript.

## Statements and Declarations

### Author contributions

TGM and RLV conceived of and designed the study with input provided by JSF. Data collection was performed by TGM with help from JSF. Data analysis was performed by TGM. The first draft of the manuscript was written by TGM with all authors contributing to the writing and editing process. All authors read and approved of the final manuscript.

### Data availability

All datasets generated during the study as well as data analysis scripts and outputs can be found on GitHub at https://github.com/tobiasgmueller/nectar_growth_assay

