## Supplemental Materials for "Nectar compounds impact bacterial and fungal growth and shift community dynamics in a nectar analog"

Yeast Media (YM)

To make 1000ml of YM, dissolve the following in deionized H_2_O

- 3g Malt Extract
- 5g Peptone
- 10g Glucose (Dextrose)
- 20g Agar
- 3g Yeast Extract

after autoclaving add

- 1mL Chloramphenicol (100 mg/mL)

Tryptone Soy Agar Media (TSA)

To make 1000ml of TSA, dissolve the following in deionized H_2_O

- 15g Tryptone
- 15g Agar
- 5g Soytone
- 5g NaCl
- 50g Fructose

after autoclaving add

- 1mL Cycloheximide (100mg/mL)

**Supplemental method 1.** The media recipes for the yeast media (YM) and tryptone soy agar (TSA) that fungi and bacteria were cultured on respectively

| **Treatment** | **Levels Found in Nectar** | **Citations** |
| --- | --- | --- |
| **4 mM H_2_O_2_** | H_2_O_2_ levels up to 4mM have been found in ornamental tobacco (*Nicotiana langsdorffii × Nicotiana sanderae*) nectar | Carter, C. et al. Tobacco Nectaries Express a Novel NADPH Oxidase Implicated in the Defense of Floral Reproductive Tissues against Microorganisms. Plant Physiol 143, 389–399 (2007); Carter, C. & Thornburg, R. W. Is the nectar redox cycle a floral defense against microbial attack? Trends in Plant Science 9, 320–324 (2004) |
| **2 mM H_2_O_2_** |  |  |
| **30% Sugar** | Sugar levels in nectar can range from 8% to over 80% | Baker, H. G. Sugar Concentrations in Nectars from Hummingbird Flowers. Biotropica 7, 37–41 (1975); Herrera, C. M., Canto, A., Pozo, M. I. & Bazaga, P. Inhospitable sweetness: nectar filtering of pollinator-borne inocula leads to impoverished, phylogenetically clustered yeast communities. Proceedings of the Royal Society B: Biological Sciences 277, 747–754 (2010) |
| **100 ng/ml Linalool** | Linalool levels can range from 5ng to over 100ng/ml in *Penstemon digitalis* nectar | Burdon, R. C. F., Junker, R. R., Scofield, D. G. & Parachnowitsch, A. L. Bacteria colonising Penstemon digitalis show volatile and tissue-specific responses to a natural concentration range of the floral volatile linalool. Chemoecology 28, 11–19 (2018) |
| **150 μg/ml BrLTP2.1 (LTP)** | Exact concentrations are unknown, however, fluorescence of BrLTP2.1 shows high levels in *brassica rapa* nectar. Previous experiments tested up to 300μg/ml | Schmitt, A. J. et al. The major nectar protein of Brassica rapa is a non-specific lipid transfer protein, BrLTP2.1, with strong antifungal activity. J Exp Bot 69, 5587–5597 (2018) |
| **22 μg/ml Deltaline** | Deltaline levels can be up to .63μg/100mg in *Delphinium* nectar, however, concentrations of the norditerpene alkaloid class as a whole can reach up to 22μg/ml in *Delphinium* nectar | Cook, D., Manson, J. S., Gardner, D. R., Welch, K. D. & Irwin, R. E. Norditerpene alkaloid concentrations in tissues and floral rewards of larkspurs and impacts on pollinators. Biochemical Systematics and Ecology 48, 123–131 (2013) |
| **1% Ethanol** | The highest reported level of ethanol in nectar is 3.8%, however, no formal survey of ethanol in floral nectar has been performed | Wiens, F. et al. Chronic intake of fermented floral nectar by wild treeshrews. PNAS 105, 10426–10431 (2008) |

**Supplemental Table 1** The concentrations of nectar compounds used as treatments along with their reported natural concentrations in floral nectar

| **Treatment** | Base nectar | 30% sugar | 2mM H_2_O_2_ | 4mM H_2_O_2_ | 100 ng/mL Linalool | 150 μg/mL LTP | .22 μg/mL Delaline | 1% Ethanol |
| --- | --- | --- | --- | --- | --- | --- | --- | --- |
| **Total made** | **25 mLs** | **25 mLs** | **25 mLs** | **25 mLs** | **25 mLs** | **15mL** | **25 mLs** | **25 mLs** |
| **g Sucrose** | 1.875 | 3.75 | 1.875 | 1.875 | 1.875 | 1.125 | 1.875 | 1.875 |
| **g Glucose** | 0.9375 | 1.875 | 0.9375 | 0.9375 | 0.9375 | 0.5625 | 0.9375 | 0.9375 |
| **g Fructose** | 0.9375 | 1.875 | 0.9375 | 0.9375 | 0.9375 | 0.5625 | 0.9375 | 0.9375 |
| **g Peptone** | 0.25 | 0.25 | 0.25 | 0.25 | 0.25 | 0.15 | 0.25 | 0.25 |
| **g Yeast Extract** | 0.75 | 0.75 | 0.75 | 0.75 | 0.75 | 0.45 | 0.75 | 0.75 |
| **mL 100x non-essential amino acids** | 12.5 | 12.5 | 12.5 | 12.5 | 12.5 | 7.5 | 12.5 | 12.5 |
| **μL 30% H2O2** | - | - | 5.67 | 11.33 | - | - | - | - |
| **μL linalool** | - | - | - | - | 2.87 | - | - | - |
| **μg LTP** | - | - | - | - | - | 2250 | - | - |
| **mg Deltaline** | - | - | - | - | - | - | 0.55 | - |
| **μL 100% Ethanol** | - | - | - | - | - | - | - | 250 |

**Supplemental Table 2** Recipes for synthetic nectar treatment solutions. All treatments were fully dissolved in deionized water before being syringe filtered through a .2μm filter to ensure sterility

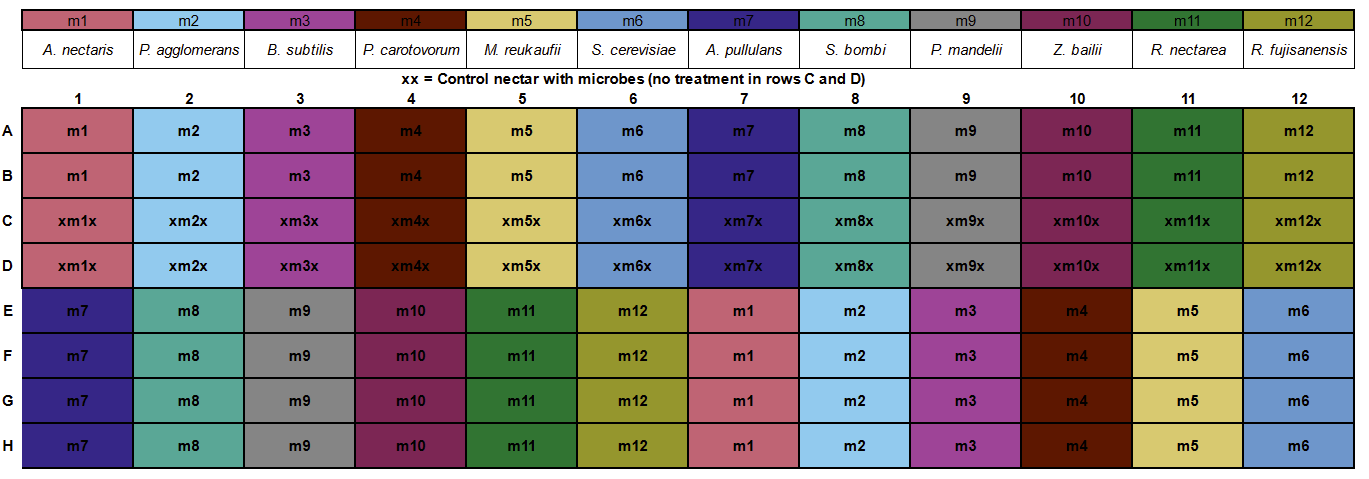

**Supplemental Figure 1** The layout of microbes on the 96 well plate. Each microbe (m1-m12, listed above) had 6 replicates in each treatment nectar (rows A:B, and E:H) and 2 replicates in control nectar (rows C:D) marked above with an X. The placement of each microbe on the plate was determined with a random number generator and kept consistent across all assays

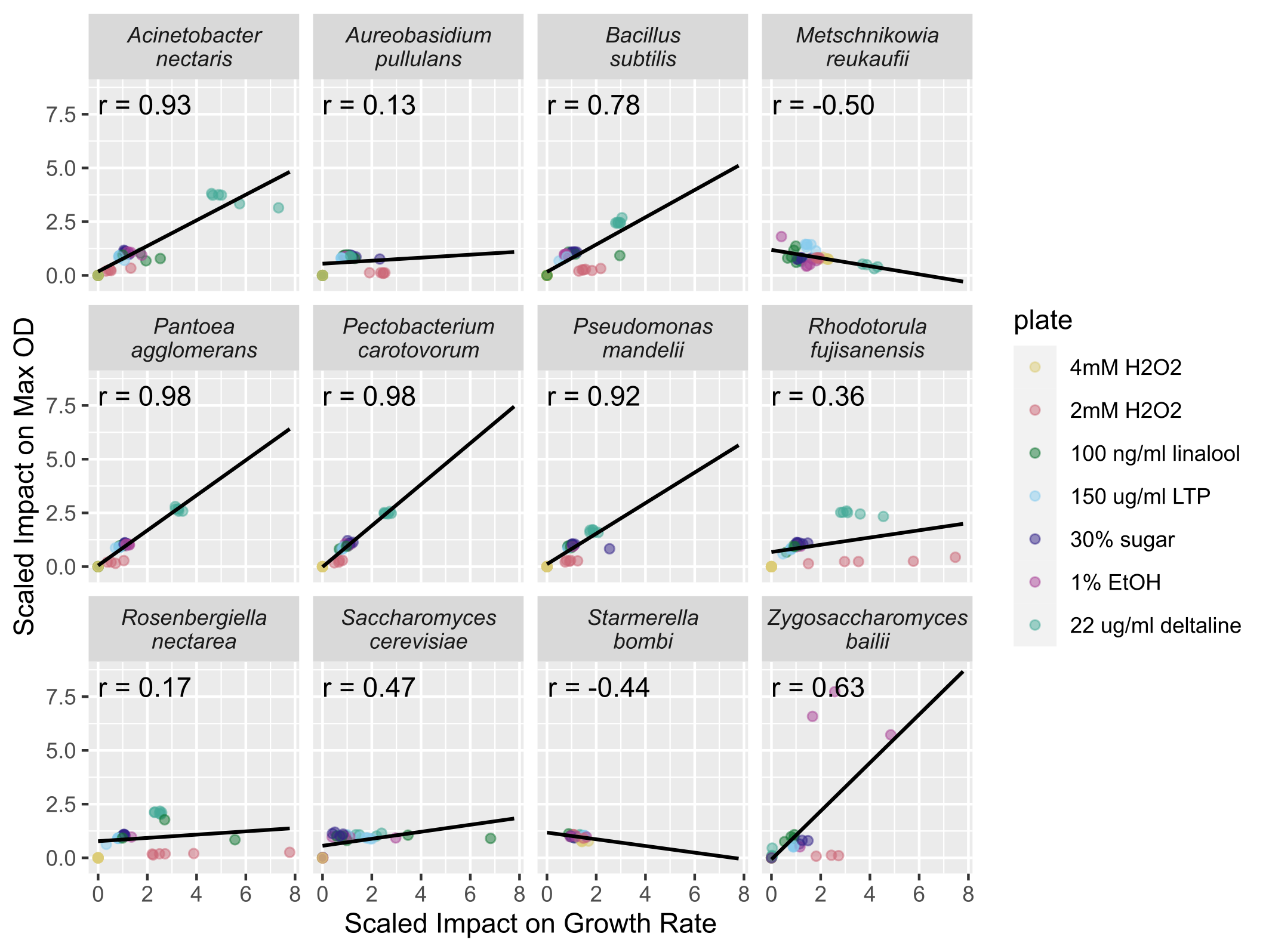

**Supplemental Figure 2** The treatment impacts on maximum OD and growth rate were correlated across many but not all species. The axes indicate the scaled effect of treatment compared to control nectar. A value of 1 represents equal max OD/growth rate in treatment and controls; values higher than one represent an increase in max OD/growth rate compared to controls and values lower than one indicate a decrease in max OD/growth rate. The Pearson's correlation coefficient (r) is given for each microbe

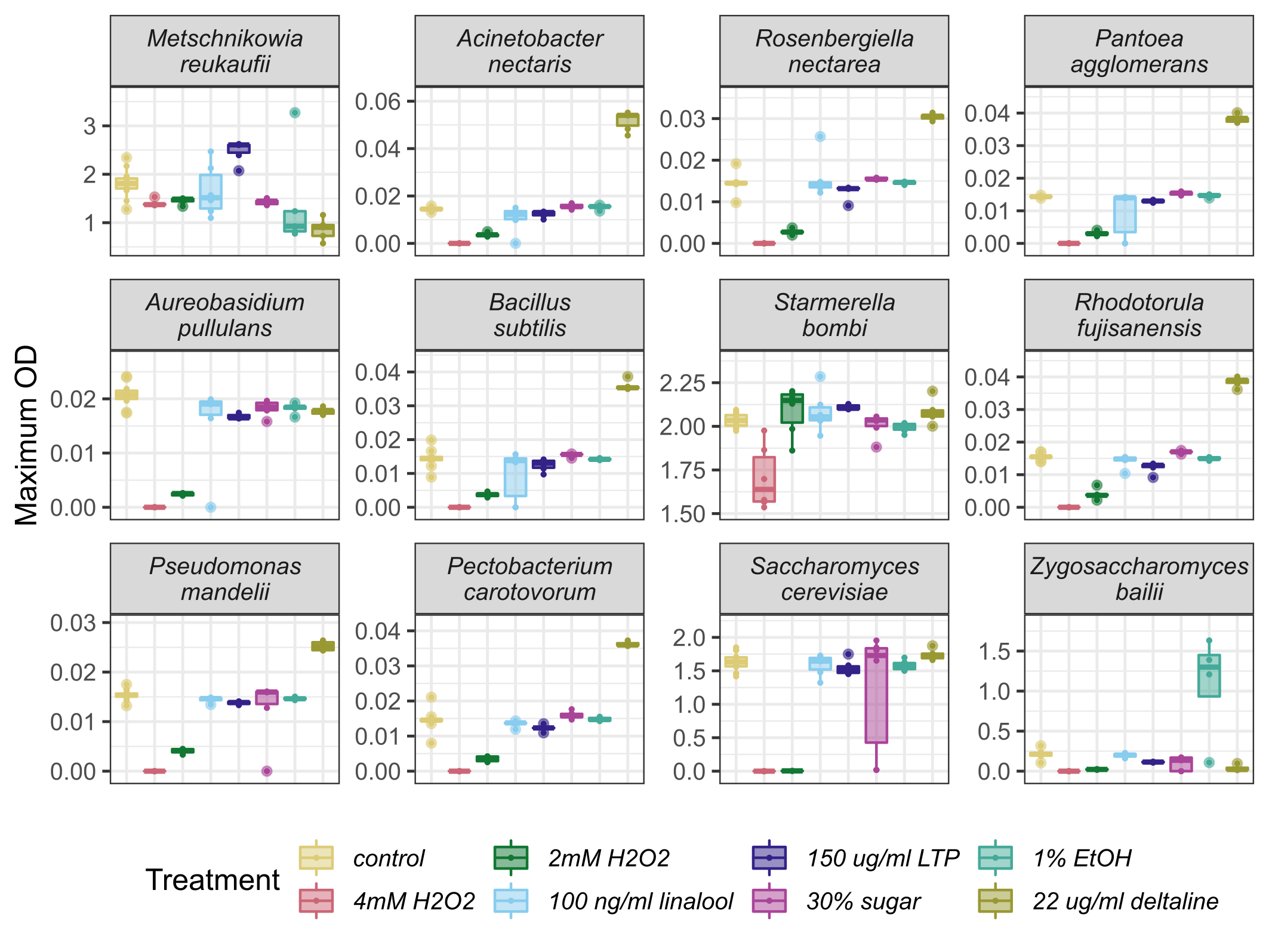

**Supplemental Figure 3.** Microbes differed in their maximum OD across different treatment nectars. Microbes are ordered from most frequently (top left) to least frequently isolated from nectar (bottom right)

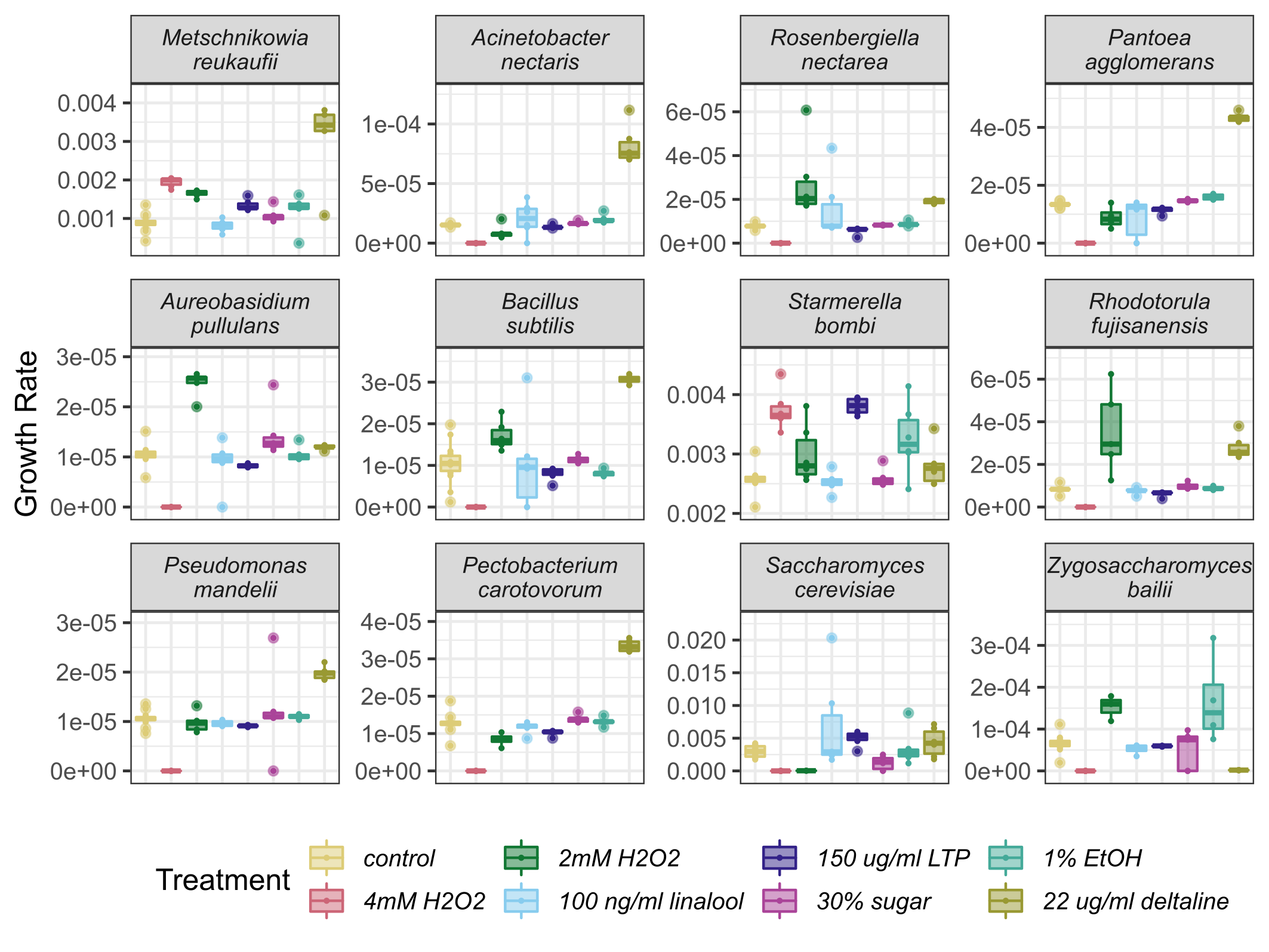

**Supplemental Figure 4.** Microbes differed in their growth rate across different treatment nectars. Microbes are ordered from most frequently (top left) to least frequently isolated from nectar (bottom right)

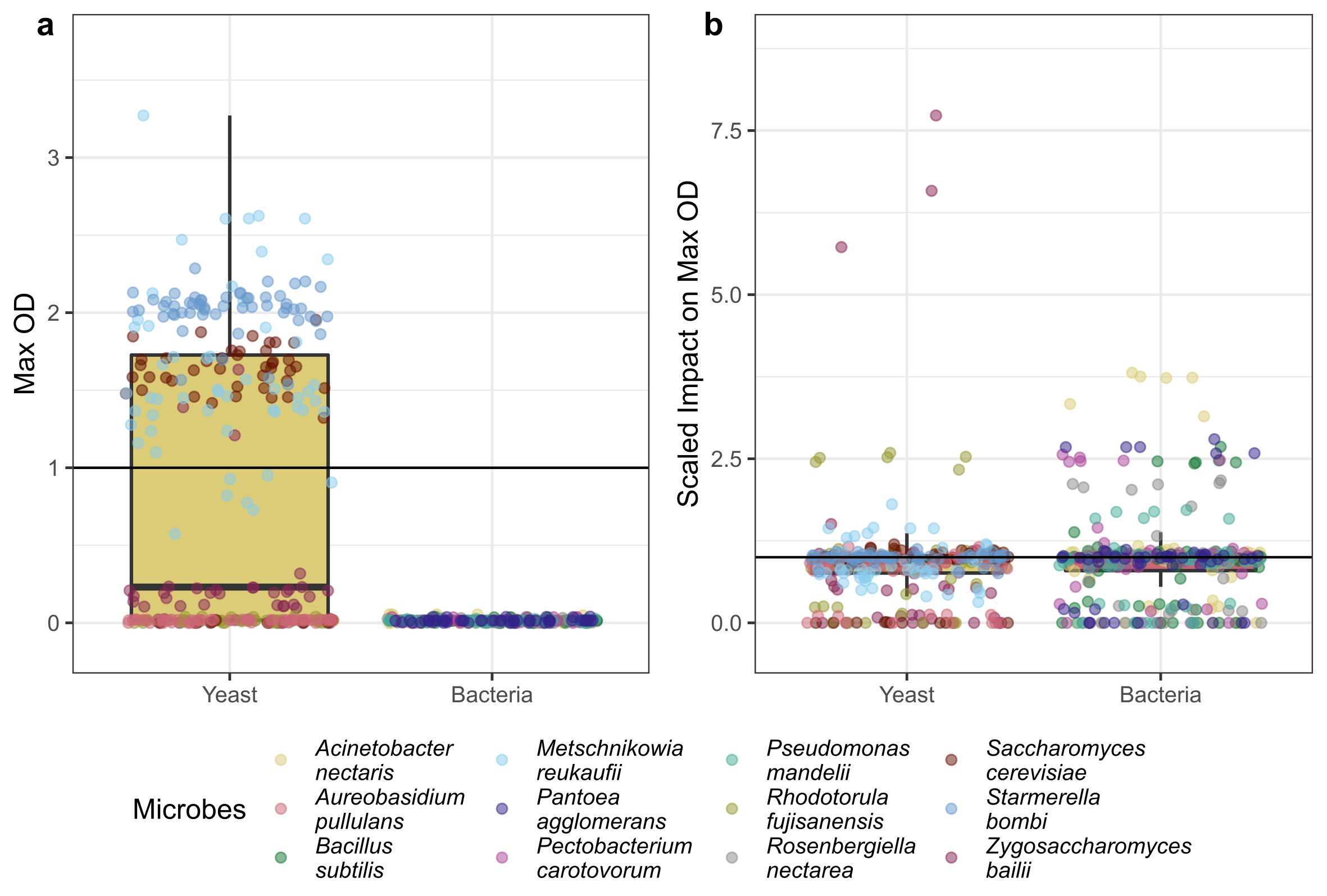

**Supplemental Figure 5.** Bacteria and Yeast differed overall in their maximum OD (panel A) but did not differ overall in their susceptibility to treatments (panel B)
